# Expression atlas of *Selaginella moellendorffii* provides insights into the evolution of vasculature, secondary metabolism and roots

**DOI:** 10.1101/744326

**Authors:** Camilla Ferrari, Devendra Shivhare, Bjoern Oest Hansen, Nikola Winter, Asher Pasha, Eddi Esteban, Nicholas J. Provart, Friedrich Kragler, Alisdair Fernie, Takayuki Tohge, Marek Mutwil

## Abstract

- The lycophyte *Selaginella moellendorffii* represents early vascular plants and is studied to understand the evolution of higher plant traits such as the vasculature, leaves, stems, roots, and secondary metabolism. However, little is known about the gene expression and transcriptional coordination of Selaginella genes, which precludes us from understanding the evolution of transcriptional programs behind these traits.
- We here present a gene expression atlas comprising all major organs, tissue types, and the diurnal gene expression profiles for *S. moellendorffii*. The atlas is part of the CoNekT-Plants database (conekt.plant.tools), which enables comparative transcriptomic analyses across two algae and seven land plants.
- We show that the transcriptional gene module responsible for the biosynthesis of lignocellulose evolved in the ancestor of vascular plants, and pinpoint the duplication and subfunctionalization events that generated multiple gene modules involved in the biosynthesis of various cell wall types. We further demonstrate how secondary metabolism is transcriptionally coordinated and integrated with other cellular pathways. Finally, we identify root-specific genes in vascular plants and show that the evolution of roots did not coincide with an increased appearance of gene families, suggesting that the existing genetic material was sufficient to generate new organs.
- Our updated database at <u>conekt.plant.tools</u> provides a unique resource to study the evolution of genes, gene families, transcriptomes, and functional gene modules in the Archaeplastida kingdom.

## Introduction

The lycophyte *Selaginella moellendorffii* is an important model organism for plant evolutionary studies and comparative genomics, as it represents lycophytes, a clade of one of the oldest still existing vascular plants on earth. It is characterised by adaxial sporangia, a root system (roots or rhizophores), a vasculature and leaf-like structures with a single vein, called microphylls. *S. moellendorffii* has a genome size of only about 100 Mbp, which is one of the smallest reported plant genomes ^1^. Lycophytes appeared about 400 Myr ago during the Silurian period and were particularly abundant during the Devonian to mid-Carboniferous period, peaking around 310 Myr ago in Euramerican coal swamps ^2^. During the Carboniferous period, many lycophytes were tall, fast-growing trees, and their remains now exist as coal. As 70% of the biomass responsible for the Bashkirian and Moscovian coal formations in Euramerica came from lycophytes ^3^, they are an important source of fossil fuels. Despite their decreasing abundance at the end of the Carboniferous period, the impact of lycophytes on the climate was significant, as their role as carbon sinks most likely contributed to the dramatic decline in atmospheric CO_2_ ^4^.

Lycophytes produce spores for reproduction that are dispersed by wind and water. One means to increase spore dispersal and outcompete neighbors in light capture, and thus facilitate the colonization of land, is to increase the height of the sporophyte. This required the evolution of specialized vasculature (xylem and phloem) for the transport of nutrients, water, and various signaling molecules and the reinforcement of cell walls by lignification ^5^. This lignification, in turn, was associated with the evolution of the phenylpropanoid metabolic pathway to synthesize and polymerize the heterogeneous aromatic lignin in the xylem cell walls ^6^.

To provide water and structural support to the larger sporophytes, land plants also evolved roots. Despite their morphological and evolutionary diversity, all roots have the capacity to acquire and transport water and nutrients, grow in a downward direction, and form branch roots, and they all possess a root cap (set of terminal protective cells), a root apical meristem (a self-sustaining stem cell population) and a radial organization of cell types (from outermost epidermis tissue to innermost vascular tissue). Root morphology in extant plants and fossil records currently suggests that roots evolved several times independently during vascular plant evolution ^5,7,8^.

In addition to the presence of vasculature and roots, land plants collectively produce more than 200,000 secondary metabolites via complex biosynthetic pathways ^9^. These secondary metabolites are not directly required for growth, reproduction, and development, but serve to provide defense against microorganisms and UV protection, and to attract pollinators. Many secondary metabolites, including those that constitute colors and flavors, are lineage-specific and play specialized roles in unique ecological niches ^10^. Selaginella contains numerous secondary metabolites, such as phenolics, alkaloids, flavonoids, lignans, selaginellins, and terpenoids ^11^. Many species of Selaginella have been used as traditional medicines for hundreds of years. In Colombia, *S. articulata* is used to treat snakebites and neutralize *Bothrops atrox* venom ^12^. In India, *S. bryopteris* is one of the plants proposed to be Sanjeevani—one that infuses life—for its medicinal properties ^13^. According to Hindu epic Ramayana, Sanjeevani was used for infusing life to Lakshmana, the younger brother of Lord Rama, who fell unconscious and nearly dead, once he was hit by a weapon during the war with a demon. While the medicinal uses of Selaginella are anecdotal, the efficacy of Selaginella metabolites has been investigated by modern approaches ^14^. For example, uncinoside A and uncinoside B biflavonoids have potent antiviral activities against respiratory syncytial virus ^15^, while biflavonoids from *S. tamariscina* inhibit the induction of nitric oxide and prostaglandins ^16,17^, which are important for the pathogenesis of some cancers ^18^. In addition, biflavone ginkgetin from *S. moellendorffii* selectively induces apoptosis in ovarian and cervical cancer cells ^19^. These findings show that the Selaginella metabolites and their biosynthetic pathways merit more investigation.

To unravel the evolution of the various organs and metabolic pathways, it is necessary to understand the functions of the underlying gene products and to compare these functions across species that possess or lack these organs and pathways ^20,21,22^. Since the experimental functional characterization of even one gene can take years, *in silico* prediction is currently the only large-scale approach to understand gene function in *S. moellendorffii*. Furthermore, classical comparative genomic approaches based solely on gene sequences are useful, but have shortcomings, as they cannot readily infer which genes work together in a pathway, i.e., form a functional gene module ^23^. Consequently, to study the evolution of new traits, we need to integrate the classical genomic approaches with predicted functional gene modules, which can be identified by expression and co-expression analysis ^24–27^. Co-expression analysis is based on the guilt-by-association principle, which states that genes with similar expression patterns across organs, developmental stages and biotic as well as abiotic stresses tend to be involved in the same or closely related biological processes ^20,21,24,28–30^. Co-expression analyses have been applied to successfully identify genes involved in cyclic electron flow ^31^, cell division ^32^, drought sensitivity and lateral root development ^29^, plant viability ^33^, seed germination ^34^, shade avoidance ^35^, and others ^20,36–43^.

To enable these comparative genomic, transcriptomic, and co-expression approaches for *S. moellendorffii*, we generated a comprehensive RNA sequencing-based (RNA-seq) expression atlas, which captures the transcriptome of the plant major organs. By uploading the expression data to CoNekT-Plants database, <u>conekt.plant.tools</u>, we provide the plant community with advanced comparative transcriptomic analyses for algae and land plants. We use the database to reveal the predicted functional gene modules involved in the biosynthesis of lignified cell walls and various secondary metabolites. We also provide examples of comparative analysis of the different cell wall types and root-specific transcriptomes. Thus, the updated CoNekT-Plants database now constitutes a unique resource to study transcriptional programs over 1 billion of years of evolution in the Archaeplastida kingdom.

## Results

### Expression atlas databases for *Selaginella moellendorffii*

The typical *S. moellendorffii* plants used in this study (Figure 1A) contained aerial root-bearing rhizophores (Figure 1B), which produce roots with the typical characteristics such as root hairs and root cap ^44^. The major photosynthetic organs of *S. moellendorffii* are the single-veined leaf-like structures called microphylls (Figure 1C), which are covered with stomata (Supplemental Figure 1A, red arrows) and contain serrated epidermal cells (Supplemental Figure 1A, green arrow). The reproductive organs are represented by sporangia-bearing structures known as strobili (Figure 1D), which are found at the tips of shoots, and contain the male micro- and the female mega-sporangia. Figures 1E and 1F show microsporangia with microspore tetrads; Supplemental Figure 1B shows strobili without sporophylls. ^1^. The microphyll structures are found along the stem, shoots (Supplemental Figure 1C) and strobili (Supplemental Figure 1D-E).

**Figure 1.**
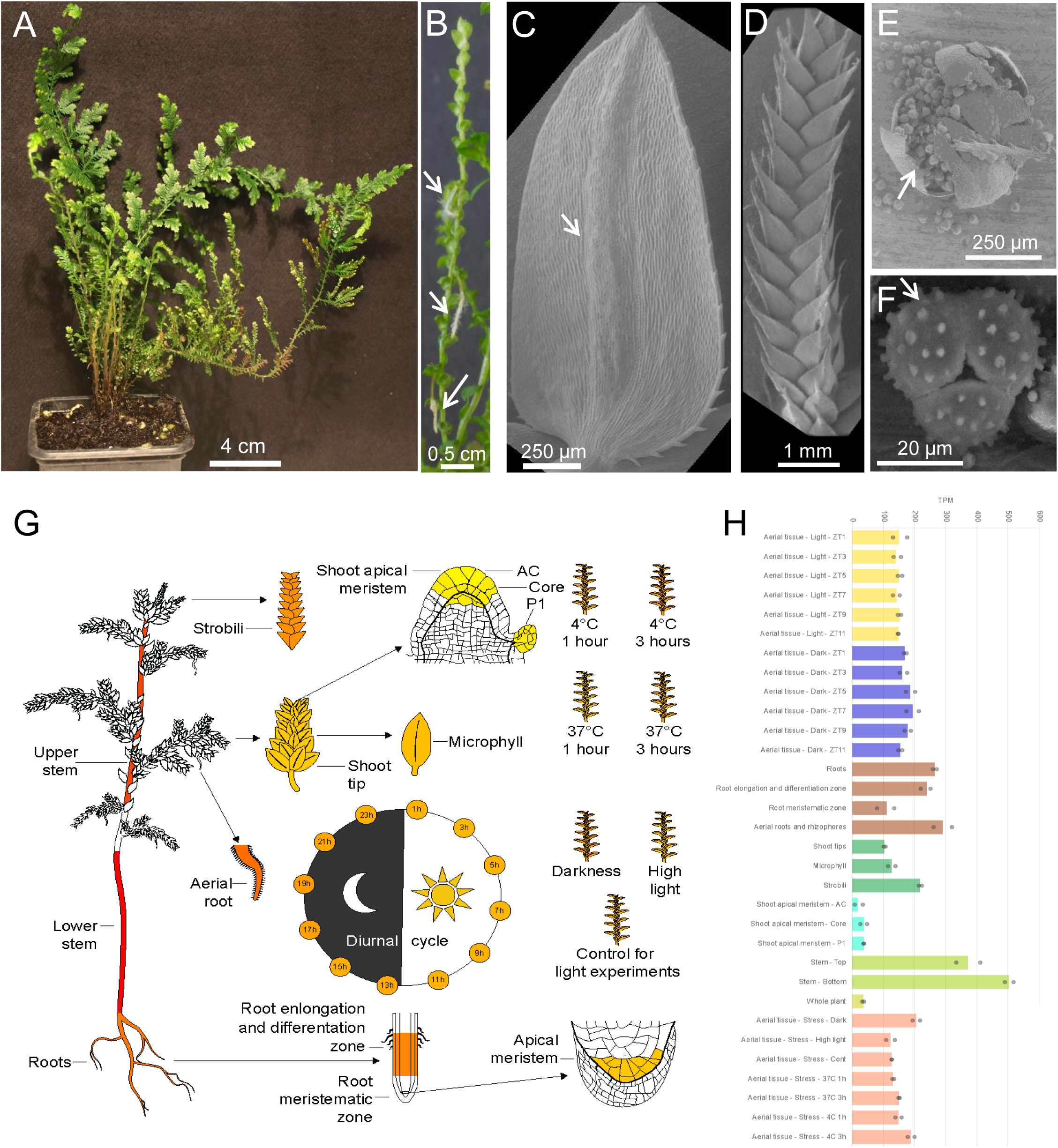
Tissues and organs used to generate the *S. moellendorffii* expression atlas. (A) A typical *S. moellendorffii* plant used in this study. (B) Stem with aerial roots (indicated by arrows). (C) to (G) Scanning electron microscope (SEM) images of Selaginella. (C) microphyll, with the vein indicated by the arrow. (D) Strobilus. (E) Crushed microsporangium with released microspore tetrads. (F) Microspore tetrad, with three microspores shown. (G) Expression profile of the *C3’H* gene from *S. moellendorffii* (*PACid_15423174*) in Selaginella eFP Browser. Expression values in the sampled organs, stress, and diurnal conditions are indicated by a color gradient. Yellow color indicates no detectable expression, while red signifies the highest expression. (H) Expression profile of the *C3’H* gene from *S. moellendorffii* (*PACid_15423174*) in CoNekT-Plants. The samples are shown on the x-axis, while the TPM expression value is indicated on the y-axis. The bars represent the average expression values and are color-coded according to the organ or experiment type. The dots indicate the minimum and maximum expression values of the replicates.

We assembled a gene expression atlas for *S. moellendorffii* by combining newly generated and publicly available RNA-seq samples from major organs, tissues, stress conditions, and a diurnal cycle study (Table 1, Supplemental Table 1, 2). The sample clustering dendrogram showed an expected clustering of replicates, similar tissues, and organs (Supplemental Figure 2). We provide two options to visualize gene expression data: (i) an *S. moellendorffii* eFP Browser (http://bar.utoronto.ca/efp_selaginella/cgi-bin/efpWeb.cgi)^45^, a graphical anatogram that color-codes the different organs and conditions based on the average expression level of a gene of interest, and (ii) CoNekT-Plants database (www.conekt.plant.tools)^46^, where the expression of each sample is shown by condition and tissue.

**Table 1.** Samples used to generate the expression atlas. Aerial roots and rhizophores: adventitious roots originated at the tips of rhizophores, strobili: the sporangia bearing structure along the stem.

| Sample description | Number of samples | Source |
| --- | --- | --- |
| Whole plant | 2 | E-MTAB-4374 |
| Roots | 2 | This study |
| Aerial roots and rhizophores | 2 | This study |
| Roots - elongation and differentiating zone | 3 | E-GEOD-64665 |
| Roots - meristematic zone | 3 | E-GEOD-64665 |
| Strobili | 2 | This study |
| Shoot tips | 2 | This study |
| Stem: top 5 cm | 2 | This study |
| Stem: bottom 5 cm | 2 | This study |
| Microphylls | 2 | This study and SRX2779511 |
| Shoot apical meristem - AC | 2 | E-MTAB-4374 |
| Shoot apical meristem - Core | 2 | E-MTAB-4374 |
| Shoot apical meristem - P1 | 2 | E-MTAB-4374 |
| 4°C: 1 hour | 2 | This study |
| 4°C: 3 hours | 2 | This study |
| 37°C: 1 hour | 2 | This study |
| 37°C: 3 hours | 2 | This study |
| Control (of dark and HL) | 1 | This study |
| Dark: 48 hours | 2 | This study |
| High light: 200 μE for 12 hours | 2 | This study |
| Diurnal expression: aerial tissue sampled every 2 hours | 39 | E-MTAB-7188 |

Since *S. moellendorffii* is a representative of early vascular plants, we present as an example the expression profile of a *p*-coumarate 3-hydroxylase (*C3’H*) gene (*Smo271465; PACid_15423174*) involved in the first steps of the lignin biosynthesis ^47^. As expected of a gene involved in vasculature biogenesis, it is highly expressed in the stem and roots indicated by the red color in the eFP Browser representation (Figure 1G) and by brown and green bars in CoNekT (Figure 1H).

### Co-expression network analysis of lignin biosynthesis reveals the association of cellulose and lignin biosynthesis

To demonstrate that our expression atlas can generate biologically meaningful co-expression networks, we investigated whether the data can produce a typical scale-free network. Scale-free topology is thought to ensure that the network remains mostly unaltered in case of disruptive mutations, and is thus an evolved property that ensures robustness against genetic and environmental perturbations ^48^. Scale-free networks show a power-law behavior where only a few genes are connected (correlated) to many genes, while the majority of genes shows only a few connections ^49^. The co-expression network of *S. moellendorffii* showed a scale-free topology, as plotting the number of connections a gene has (node degree) against the frequency of this association produced a negative slope (Figure 2A), confirming the scale-free topology of our network and suggesting the biological validity of our expression data.

**Figure 2.**
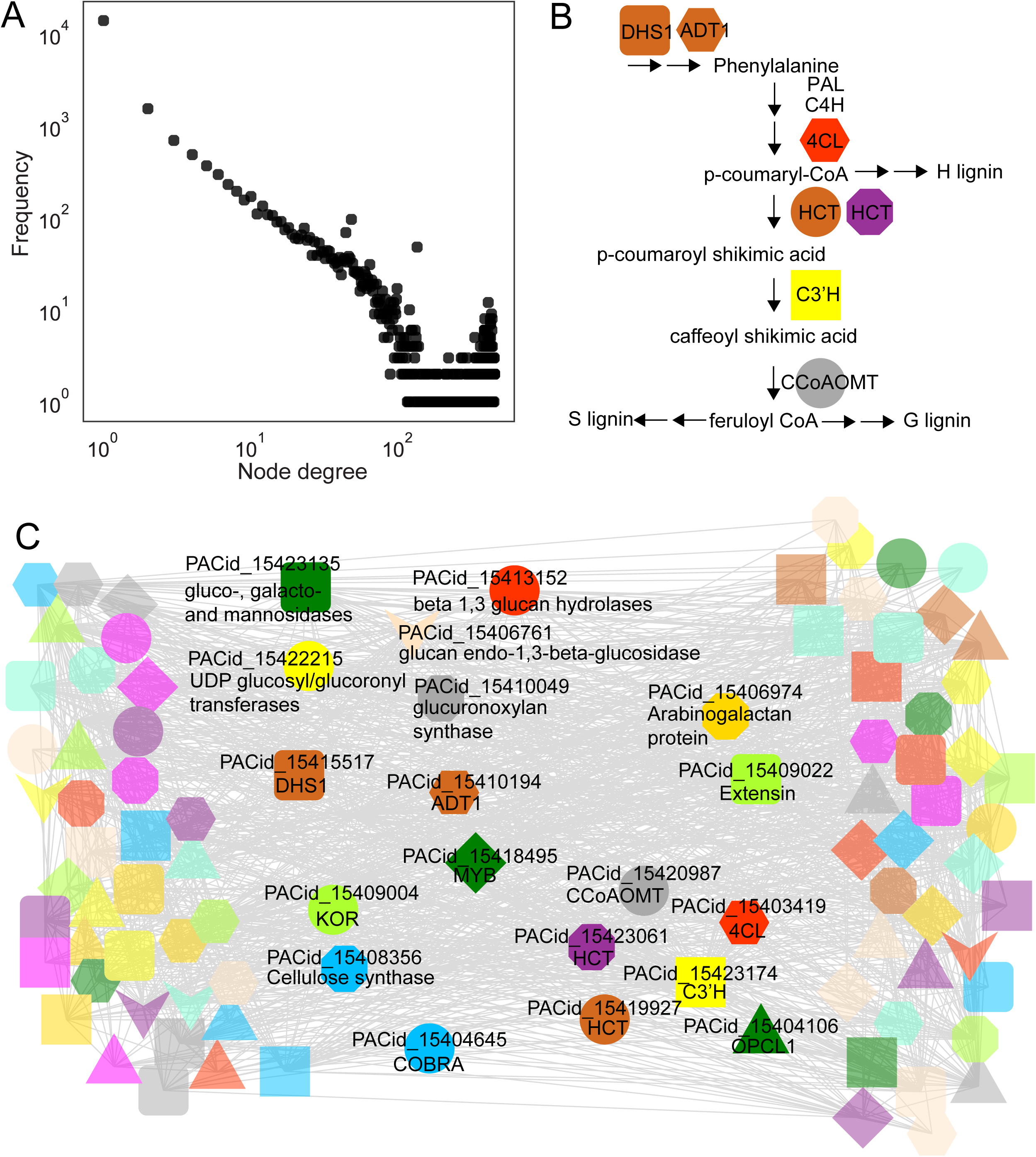
Co-expression network and CoNekT-Plants instance of *S. moellendorffii*. (A) Power-law plot obtained from *S. moellendorffii* expression data. The x-axis shows the node degree (number of co-expression connections of a gene), while the y-axis indicates the frequency of a degree. The two axes are log_10_-transformed. (B) Simplified pathway of lignin biosynthesis. Colored nodes indicate orthogroups of the involved enzymes. Enzymes without the colored nodes, such as *PAL* and *C4H*, indicate the enzymes not found in the *C3’H* module. (C) Co-expression network of *S. moellendorffii C3’H* gene *PACid_15423174* (yellow square). Nodes indicate genes, edges connect co-expressed genes. Node colors/shapes indicate which genes belong to the same orthogroup.

Since lignocellulosic cell walls are the main characteristics of the plant vasculature, we decided to investigate the biosynthetic module of lignin in Selaginella. Lignin is produced by the phenylalanine/tyrosine pathway through the formation of lignin monomers followed by their polymerization in the cell wall ^50^. The three monolignols which are incorporated into the lignin polymer are guaiacyl (G), syringyl (S), and *p*-hydroxyphenyl (H) ^51^ which polymerise into G-lignin, S-lignin and H-lignin, respectively (Figure 2B). Although the appearance of S-lignin was thought to be specific to angiosperms due to its absence in conifers and ferns, S-lignin is found in Selaginella ^52^, suggesting the independent evolution of S-lignin in lycophytes and angiosperms.

The co-expression network neighborhood of the *C3’H* gene (*PACid_15423174*) contained genes encoding enzymes involved in the early stages of lignin biosynthesis (Figure 2C)^53^. The enzymes are 4-coumarate-CoA ligase (4CL, red hexagon), hydroxycinnamoyl-CoA shikimate/quinatehydroxycinnamoyl transferase (HCT, represented by two genes in two different orthogroups) and caffeoyl-CoA *O*- methyltransferase (CCoAMT, gray circle) (Figure 2C). We also observed genes for two enzymes involved in the biosynthesis of phenylalanine DHS1 (3-DEOXY-D-ARABINO-HEPTULOSONATE 7- PHOSPHATE SYNTHASE 1)^51^ and ADT (AROGENATE DEHYDRATASE 1)^54^, suggesting that biosynthesis of lignin and phenylalanine are coupled, probably to coordinate the high demand of this amino acid in lignin biosynthesis. In addition, we identified key genes of cell wall biosynthesis such as cellulose synthase, COBRA, and KORRIGAN (KOR)^55,56^, various cell wall proteins (arabinogalactan protein, extensin) and various polysaccharide-acting enzymes. Furthermore, we observed a member of the MYB transcription factor family that is, among other pathways, known to regulate vasculature formation (Figure 2C, green diamond)^57^. Importantly, similar modules containing genes involved in lignin, phenylalanine, cell wall biosynthesis, and MYB regulators have been observed in *Arabidopsis thaliana* and *Brachypodium distachyon* ^41^ suggesting that the conserved module shown in Figure 2C indeed evolved in early vascular plants and is thus at least 400 million years old.

### Expression and phylogenetic analyses reveal the duplication and sub-functionalization of cell wall biosynthesis modules

The composition of cell walls evolved to accommodate for the different niches of land plants, with variations not only across different species but also among different organs within the same species ^58^. *Arabidopsis thaliana* contains at least four cell wall types: (i) the ubiquitous primary cell wall (PCW), (ii) the secondary cell wall present in the vasculature (SCW), and the tip growing cell walls of (iii) root hairs and (iv) pollen tubes ^59^. Since *S. moellendorffii* contains PCW, bifurcating roots with root hairs (Figure 3A)^60^ and vasculature with SCW (Figure 3B), it provides a unique reference to study the evolution of these four types. To this end, we analyzed the phylogenetic trees and gene expression of the six major gene families involved in cellulosic cell wall biosynthesis: (i) cellulose synthases (*CesA*) that have subfunctionalized to PCW and SCW (Figure 3C)^56^, (ii) cellulose synthase-like D genes (*CslD*) involved in tip-growth (Figure 3D)^61^, (iii) *CHITINASE-LIKE* (*CTL*, Figure 3E), (iv) *KORRIGAN* (*KOR*, Figure 3F), (v) *COMPANION OF CELLULOSE SYNTHASE* (*CC*, Figure 3G), and (vi) *COBRA* (*COB*).

**Figure 3.**
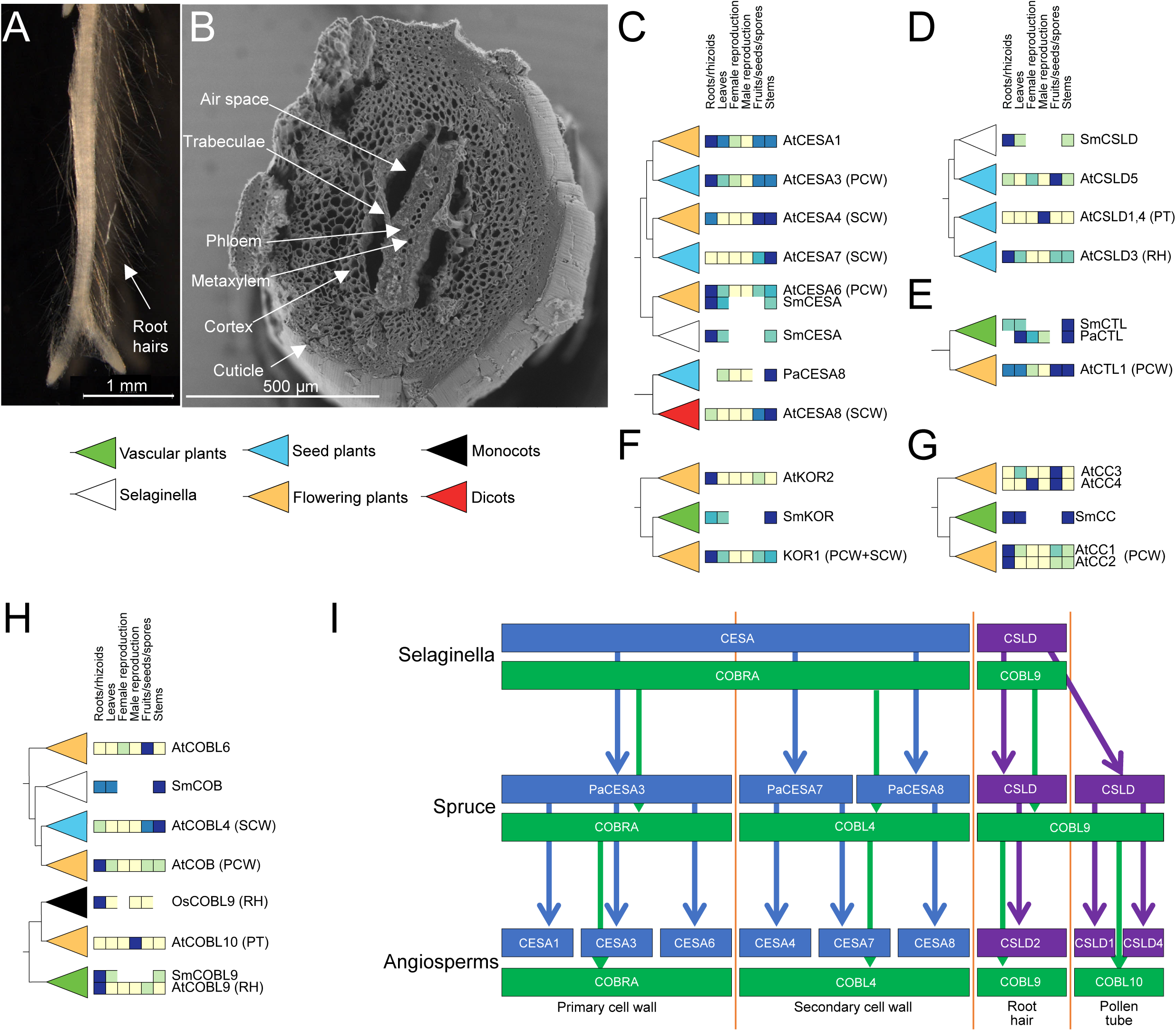
Phylogenetic and expression analysis of cell wall biosynthesis genes. (A) *S. moellendorffii* bifurcating root tip, with root hairs. (B) SEM image of *S. moellendorffii* stem, with the major tissue types indicated. Phylogenetic trees from CoNekT-Plants for (C) cellulose synthases (*CesAs*), (D) cellulose synthase-like D (*CslDs*), (E) chitinase-like (*CTL*), (F) korrigan (*KOR*), (G) companion of cellulose synthase (*CC*) and (H) *COBRA*. For brevity, only the representative genes from Arabidopsis (genes starting with At), *S. moellendorffii* (Sm), spruce (Pa) and rice (Os) are shown. The color of the leaves indicates the oldest lineage found in the clade. For example, the red color indicates that only dicot genes are found in a clade. The colored boxes to the left of the gene identifiers correspond to the expression in major organs, where light color indicates low expression, while dark color shows high gene expression. Genes involved in primary cell wall (PCW), secondary cell wall (SCW), root hairs (RH) and pollen tubes (PT) are indicated. (I) Proposed model of the evolution of genes involved in PCW, SCW, RH, and PT. The duplication and sub-specialization events are shown for *CesAs* (blue boxes), *COBRA* (green) and *CslDs* (purple).

The CesA family contains multiple non-redundant copies in angiosperms, where PCW *CesAs* are typically ubiquitously expressed, while SCW *CesAs* show specific expression in roots and stems ^59^. We observed that *S. moellendorffii CesA* genes (Figure 3C, Supplemental Figure 3) are all found on the same clade as PCW *AtCesA6* and are absent from any SCW clades. This suggests that *S. moellendorffii CesAs* are part of one module that produces both PCW and SCW. Conversely, in seed plants, the *CesAs* duplicated and subfunctionalized to produce PCW (*CesA3*) and SCW (*CesA7* and *8*), and in flowering plants, *CesA3* further divided into *CesA1, 3, 6*, while SCW *CesA7* duplicated to produce *CesA4* and *7* (Figure 3C, Supplemental Figure 3).

The *CslD* family is essential for tip growth of root hairs and pollen tubes, and we observed one copy of *CslD* in *S. moellendorffii* expressed in roots (Figure 3D, Supplemental Figure 4), suggesting its function in root hair growth (Figure 3A). Conversely, in seed plants, three clades of *CslD* genes were formed, where each clade shows specific expression in roots (root hair *CslD3* clade), the male gametophyte (pollen tube *CslD1* clade) or a more ubiquitous expression (cell plate *CslD5* clade). In the ancestor of angiosperms, the pollen tube *CslDs* further duplicated and subfunctionalized into non-redundant *CslD1* and *4* (Figure 3D, Supplemental Figure 4).

While we did not observe any evidence for subfunctionalization for *CTLs* (Figure 3E, Supplemental Figure 5), *KORs* (Figure 3F, Supplemental Figure 6) or *CCs* (Figure 3G, Supplemental Figure 7), the COBRA family showed distinctive grouping with PCW and SCW genes in flowering plants (Figure 3H, Supplemental Figure 8). Conversely, in *S. moellendorffii* only one ubiquitously expression clade was associated with the seed plant PCW and SCW clades, suggesting that *S. moellendorffii COBRA* is involved in the biosynthesis of both cell wall types. Interestingly, another *COBRA* from *S. moellendorffii* was found on the same clade as root hair *COBL9* from Arabidopsis (Figure 3H, Supplemental Figure 8), suggesting that the subfunctionalization of the COBRA family into tip growing and non-tip growing cell walls took place in the ancestor of vascular plants. Tip growing *COBL9* further subfunctionalized into pollen tubes and root hairs in the ancestor of angiosperms (Figure 3H, Supplemental Figure 8).

Taken together, based on the phylogenetic trees and the expression profiles that we obtained from CoNekT-Plants, we propose a model describing the duplication and subfunctionalization events of the cell wall modules (Figure 3I). While *S. moellendorffii* contains both PCWs and SCWs, these two cell wall types are biosynthesized by one type of *CesAs* and *COBRAs*. The PCW/SCW subfunctionalization of *CesAs* and *COBRAs* took place in the ancestor of seed plants, and the *CesAs* further subfunctionalized into at least three non-redundant isoforms (PCW: *CesA1, 3, 6* and SCW: *CesA4, 7, 8*) in the ancestor of flowering plants. Since *S. moellendorffii* only contains root hairs, only one clade of *CslDs* and *COBL9* are found in lycophytes, but *CslDs* duplicated and subfunctionalized to support pollen tubes that arose in the ancestor of seed plants. In the ancestor of angiosperms, the *COBL9* further subfunctionalized into root hair- and pollen tube-specific forms, while the pollen tube *CslD* gene subfunctionalized into the non-redundant *CslD1* and *CslD4*.

### Analysis of functional modules in *S. moellendorffii* and the Archaeplastida kingdom

Co-expression networks contain clusters of highly connected genes (modules), which represent functionally related genes ^20,33,62^. We used the Heuristic Cluster Chiseling Algorithm (HCCA)^33^ to identify 365 clusters of co-expressed genes in *S. moellendorffii* (Supplemental Table S3), and investigated their biological functions by enrichment analysis of the MapMan functional bins. We found that 155 clusters were significantly enriched (FDR adjusted p-value < 0.05) for at least one biological process (Figure 4, Supplemental Figure 9). All the biological processes captured by MapMan were enriched in at least one cluster, with the exception of ‘S-assimilation’ (Figure 4, Supplemental Figure 9). Most of the processes, such as ‘photosynthesis’, ‘cell wall’ (comprising various cell wall-acting pathways), ‘not assigned’ (unknown), ‘protein’ (biosynthesis, degradation and modification), ‘RNA’ (regulation of transcription), ‘misc’ (various enzymes) and others, were enriched in more than one cluster suggesting that multiple functional modules are involved in these processes. Moreover, various processes were found in the same cluster, e.g., ‘cell wall’, ‘amino acid metabolism’, ‘secondary metabolism’ and ‘C1 metabolism’ were present in the same cluster (Figure 4A, Supplemental Figure 9, blue rectangle). This is in line with the observed coordination of lignin and cell wall biosynthesis (Figure 2)^63^, and suggests a transcriptional coupling of these and other processes.

**Figure 4.**
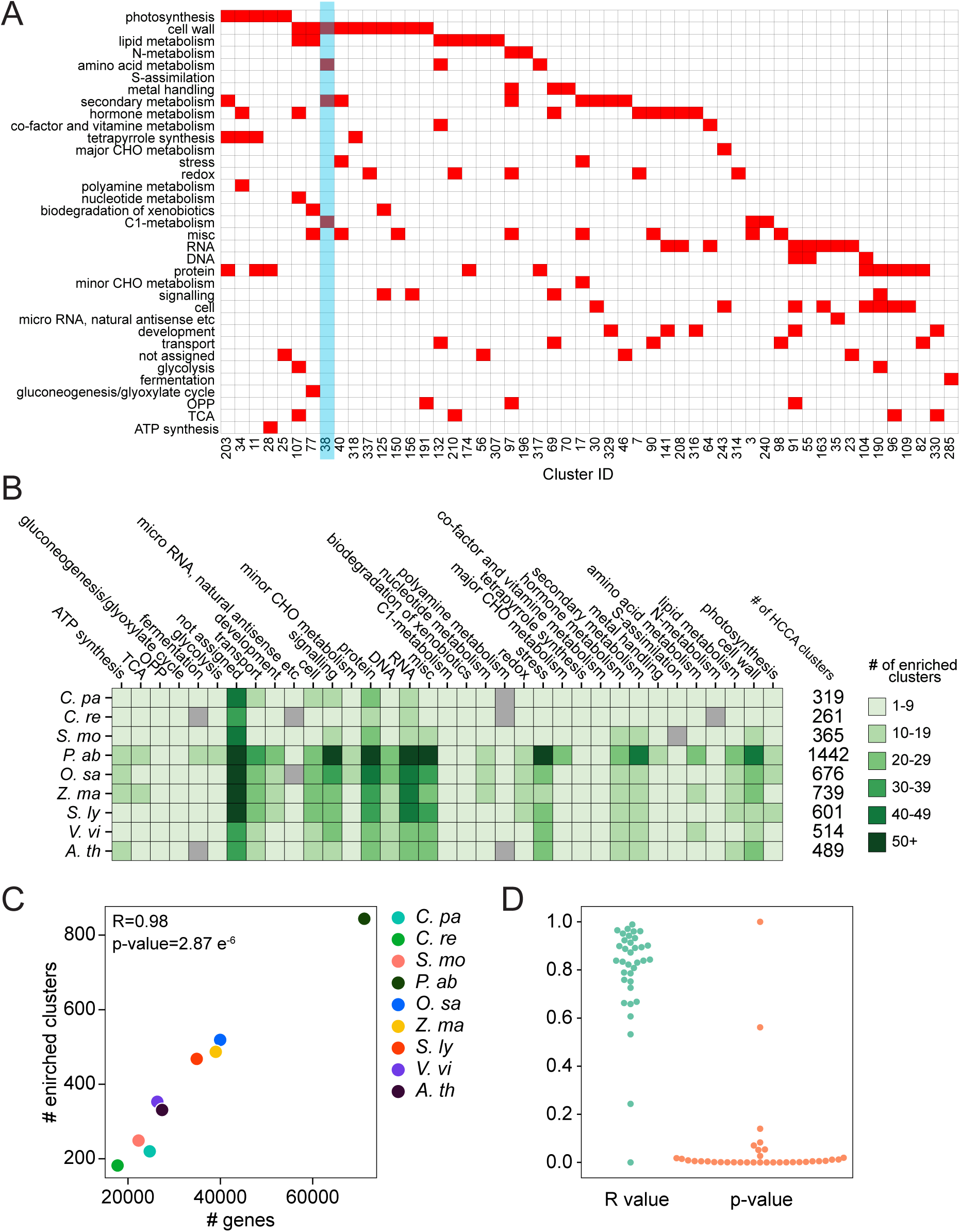
Comparative analyses of biological functions of the co-expression clusters. (A) Heuristic Cluster Chiseling Algorithm (HCCA) clusters of *S. moellendorffii*. The columns show cluster ID, while the rows correspond to MapMan bins, that represent biological processes. Clusters enriched for a MapMan bin (FDR adjusted p-value < 0.05), are indicated by red cells. Only 52 out of 365 clusters are shown in this figure, for brevity. (B) The number of enriched bins found in the HCCA clusters of CoNekT-Plants species. The species are shown in rows, while the bins are indicated in the columns. The six shades of green indicate the number of clusters assigned to a bin. (C) Relationship between the number of genes (y-axis) and the number of HCCA clusters enriched for at least one MapMan bin (y-axis), for each of the nine species in the CoNekT database (indicated by colored points). The Pearson r-value and the resulting p-value are indicated. (D) Correlation between the number of genes assigned to a bin and the number of clusters enriched for the bin. Each point represents a MapMan bin. Blue points correspond to R-values, while the orange points correspond to the p-values.

To investigate whether *S. moellendorffii* shows a typical number of functional modules, we plotted the number of functionally enriched clusters for the other species in CoNekT-Plants, which contains two algae (*Cyanophora paradoxa, Chlamydomonas reinhardtii*) and seven land plants (*Selaginella moellendorffii, Picea abies, Oryza sativa, Zea mays, Solanum lycopersicum, Vitis vinifera, Arabidopsis thaliana*). Despite a marked increase in the number of clusters in seed plants, the nine species show a similar distribution of enriched clusters for most processes (Figure 4B). The vascular plant-specific expansion includes ‘misc’ (miscellaneous, containing various enzymes), ‘hormone metabolism’, ‘secondary metabolism’ and ‘cell wall’. This is in line with evolutionary studies which revealed that hormone metabolism and secondary metabolism appeared to provide chemical defense and response to dehydration when plants started to colonize the land ^1,64,65^.

The increased number of clusters can be explained by a simple linear relationship between the number of genes and the enriched clusters belonging to any biological process. For example, the number of genes in a species positively correlates with the genome-wide number of clusters enriched for any biological process (Figure 4C). This positive correlation is seen for almost all biological processes, e.g., the number of genes assigned to a cell wall bin is strongly correlated to the number of its enriched clusters. The few exceptions to this rule (7 out of 35 bins) are involved in various aspects of primary metabolism (Figure 4D, Supplemental Table 4). These results demonstrate that the increased number of genes results in new co-expression clusters, possibly harboring novel functions.

### Secondary metabolic pathways are co-regulated with specific biological processes

When plants colonized land, new organs, biosynthetic pathways, and compounds evolved to enable plants to adapt to the new environment. These innovations comprised novel tissues and organs such as the vasculature, roots, and flowers, and various mechanisms to resist desiccation, to establish a sophisticated intracellular communication and to interact with other organisms for defense and reproduction ^9,66–69^. The latter innovations required the establishment and expansion of gene families involved in hormone biosynthesis and secondary metabolism (Figure 4B)^1,65^.

We first analyzed the number of gene families which take part in the various secondary metabolic pathways (Figure 5A). As previously observed ^65^, we noticed a ∼5-fold increase in the number and diversity of families associated with secondary metabolism in vascular plants compared to algae (Figure 5A). Except for spruce (*P. abies*), which showed an exceptionally high number of gene families (663) associated with secondary metabolism, all vascular plants showed ∼200 gene families involved in secondary metabolism (Figure 5A). The higher number of secondary metabolite gene families in spruce can be explained by a more than two-fold higher number of total genes compared to Arabidopsis (71,158 genes vs. 27,382 genes, respectively).

**Figure 5.**
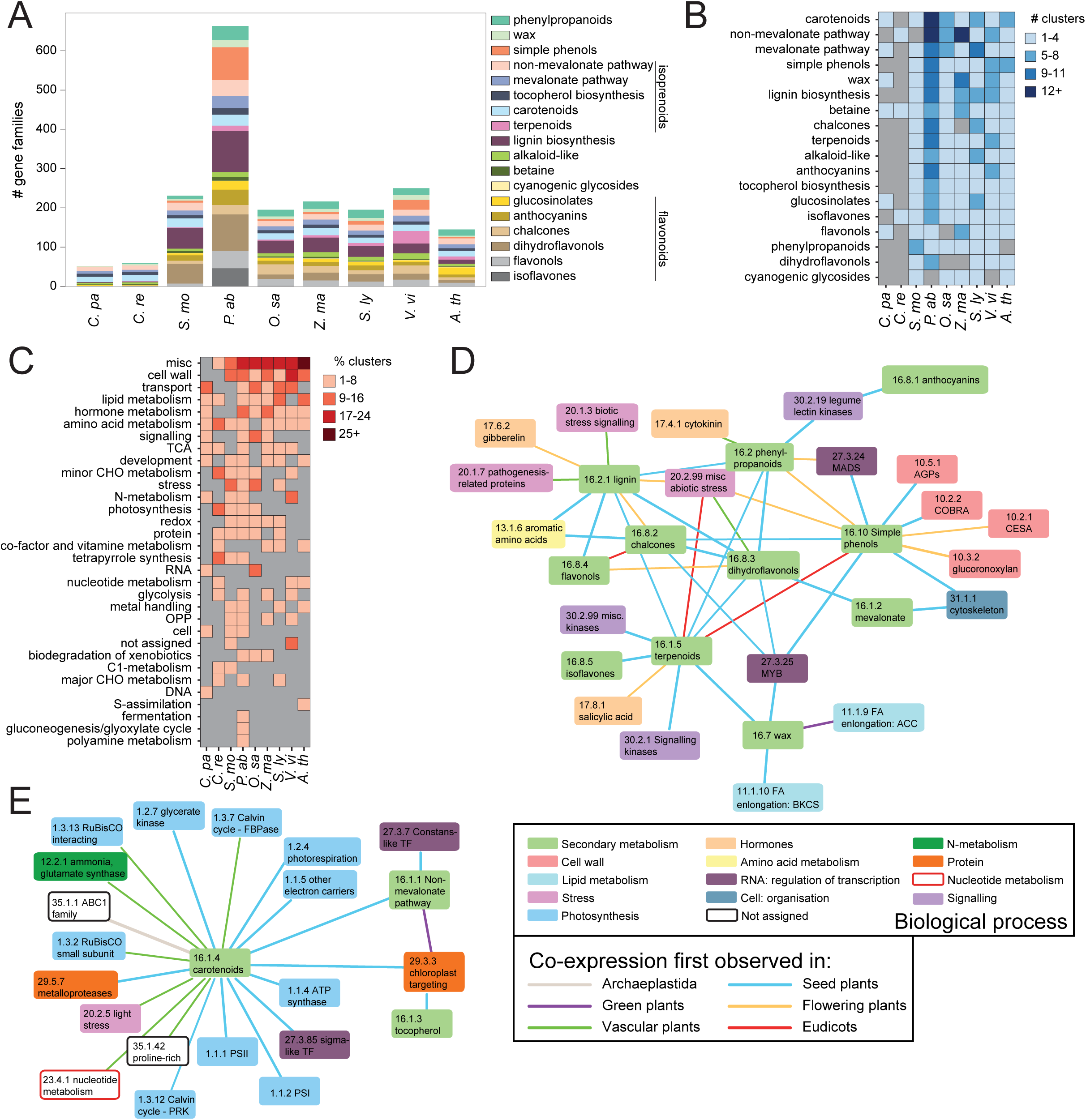
Analysis of co-expression patterns of clusters involved in secondary metabolism. (A) The number of gene families (y-axis) assigned to the different secondary metabolite MapMan bins in the species found in CoNekT-Plants (x-axis). (B) The number of HCCA clusters enriched for secondary metabolite bins, indicated by four shades of blue. Gray color (e.g., terpenoids in algae) indicates that no clusters for a specific bin in a species were found. (C) Co-occurrence analysis of secondary metabolite and other bins. The percentage of non-secondary metabolite bins enriched in clusters enriched for secondary metabolite bins. The four shades of burgundy indicate the percentage, while gray cells show that a given bin is not enriched with secondary metabolite bins. (D) Bin co-occurrence network. Nodes represent MapMan bins color-coded according to the type, while edges connect bins that were enriched in the same cluster in at least three species. The edges indicate the oldest phylostratum where the bins co-occurred in a cluster. (E) Bin co-occurrence network for the photosynthesis-related processes.

To identify the secondary biosynthetic modules in the Archaeplastida kingdom, we calculated which bins belonging to the ‘secondary metabolism’ are enriched in the HCCA clusters (Figure 5B). While only a few clusters were enriched for any secondary bins in the algae *C. paradoxa* and *C. reinhardtii*, vascular plants contained clusters enriched for nearly all secondary metabolites (Figure 5B). In line with the observation that the number of genes is correlated with the number of enriched clusters, spruce showed the highest amount of clusters enriched for secondary metabolism (Figure 5B). Interestingly, we observed a species-specific expansion of distinct secondary metabolite pathways, such as lignin biosynthesis (phenylpropanoids) and dihydroflavonols in *S. moellendorffii*, terpenoids in *V. vinifera* and glucosinolates in *A. thaliana* (Figure 5B, Supplemental Figure 10). Spruce showed a distinct expansion for carotenoid and non-mevalonate pathway clusters, which can be attributed to the high amounts of photoprotective xanthophylls produced in needles and the high emissions of isoprenoids from spruce ^70^.

Next, we investigated how diverse biological processes are co-expressed with secondary metabolism. As exemplified by lignin and cellulose biosynthesis (Figure 2C), functionally related processes can be transcriptionally coordinated (Figure 4A)^71^. To explore how the secondary metabolism is integrated with other biological pathways, we investigated which MapMan bins tend to be co-enriched with the secondary metabolism bin in the HCCA clusters. Figure 5C shows that the bin ‘misc’ was most frequently co-enriched with secondary metabolism. This bin includes several enzymes necessary for secondary metabolite biosynthesis, such as glutathione S transferases for the biosynthesis of sulfur-containing secondary metabolites ^72^, cytochrome P450 for lignin biosynthesis ^51^ and O-methyltransferases involved in the biosynthesis of phenylpropanoids, flavonoids, and alkaloids ^73^. The other five most frequently co-enriched bins are ‘cell wall’, ‘transport’, ‘lipid metabolism’, ‘hormone metabolism’ and ‘amino acid metabolism’ (Figure 5C), suggesting that secondary metabolic pathways are tightly associated with these processes.

To reveal biologically relevant associations of secondary metabolism with the processes in Figure 5C, we created a network, where the nodes represent processes and the edges connect processes present in the same cluster in at least 3 species (Figure 5D, Supplemental Table 5 shows the association of all processes in all species). The color of the edges indicates when the two processes were first observed to be associated. Most of the associations appeared in seed plants (Figure 5D-E), which can be explained by a large number of HCCA clusters found in spruce (Figure 5A, B). The network shows that ‘cell wall’ is transcriptionally associated with simple phenols, the building blocks of lignin. Lignin is in turn connected to stress responses, which is in line with the observed lignin deposition changes upon biotic and abiotic stresses ^74^, such as cold ^75^, drought stress ^76^, and mechanical stress ^77^. Interestingly, secondary metabolite bins such as simple phenols, chalcones, dihydroflavonons and phenylpropanoids were co-enriched with MYB and MADS transcription factors (Figure 5D), which is in line with the known regulatory role of these transcription factors in these processes ^78^. Moreover, we observed a large association between carotenoids and the photosynthesis and carbon fixation machinery (Figure 5E), which is in agreement with the role of these pigments in promoting photosynthesis and photoprotection ^79,80^. The carotenoid bin was also associated with sigma-like transcription factors which regulate expression of photosynthesis genes ^81^. These and numerous other associations demonstrate the biologically meaningful integration of secondary metabolism with the cellular machinery, and can be used to further understand the wiring of biological processes in response to the environment.

### Expression and phylostratigraphic analyses reveal a highly conserved root transcriptome

Roots have evolved to ensure anchorage to the soil and to allow efficient water and nutrient acquisition in land plants. In contrast to the tip-growing rhizoids in early, non-vascular land plants, all vascular plants possess roots that include a root cap, an apical meristem and a radial cell type organization ^5^. Despite the differences observed between vascular plants regarding root morphology and architecture, the root transcriptomes are highly conserved, which suggests a unique, or highly convergent, origin of roots ^82^.

To further study the evolution of roots, we analyzed root transcriptomes by using the “specific profile” feature in CoNekT-Plants, which can identify genes specifically expressed in organs and conditions (https://conekt.sbs.ntu.edu.sg/search/specific/profiles). To detect expression specificity, CoNekT uses a specificity measure (SPM) that ranges between zero (gene is not expressed in a given sample) to one (gene is exclusively expressed in the sample)^83^. By using SPM>0.85 cutoff and selecting ‘Roots/rhizoid’ as target organ, we obtained 4314, 12177, 4316, 4185 and 3247 genes for *S. moellendorffii, O. sativa, Z. mays, S. lycopersicum*, and *A. thaliana*, respectively (Supplemental Table 6). As CoNekT-Plants does not contain root samples for *P. abies* and *V. vinifera*, they were omitted from the analysis. Since roots first appeared in the ancestor of vascular plants, we hypothesized that the root-specific gene families also emerged in this period. To investigate this, we applied a phylostratigraphic analysis, where the age of a gene family is based on its presence in the most basal, i.e. earliest, plant ^24^. The different phylostrata are “Archaeplastida”, “Green plants”, “Vascular plants”, “Seed plants”, “Flowering plants”, “Monocots”, “Eudicots” and “Species-specific”. For example, a gene family containing *S. moellendorffii* genes and Arabidopsis genes would be assigned to the “Vascular plants” phylostratum, while a family containing *S. lycopersicum* (tomato) genes and Arabidopsis genes would be assigned to the “Eudicot” phylostratum.

Our phylostratigraphic analysis revealed a quite similar distribution of the different phylostrata in the root-specific transcriptomes between the five different species (Figure 6A). The following enrichment analysis showed that the “Vascular plants” phylostratum gene families were not over-represented in the roots (Figure 6A, green bars, Supplemental Table 7). Conversely, the “Species-specific” phylostratum, which represents gene families that are only found in one species (Supplemental Table 6), was significantly over-represented in most of the plants (Figure 6A, dark-orange bars, Supplemental Table 7). These results suggest that the evolution of the root organ in the ancestor of vascular plants did not require the acquisition of novel genetic material. Conversely, the significant enrichment (FDR adjusted p-value < 0.05) of gene families of the “Species-specific” phylostratum suggests that the different root morphologies observed in these plants required the generation of novel functions based on new genetic material (Figure 6A)^82^.

**Figure 6.**
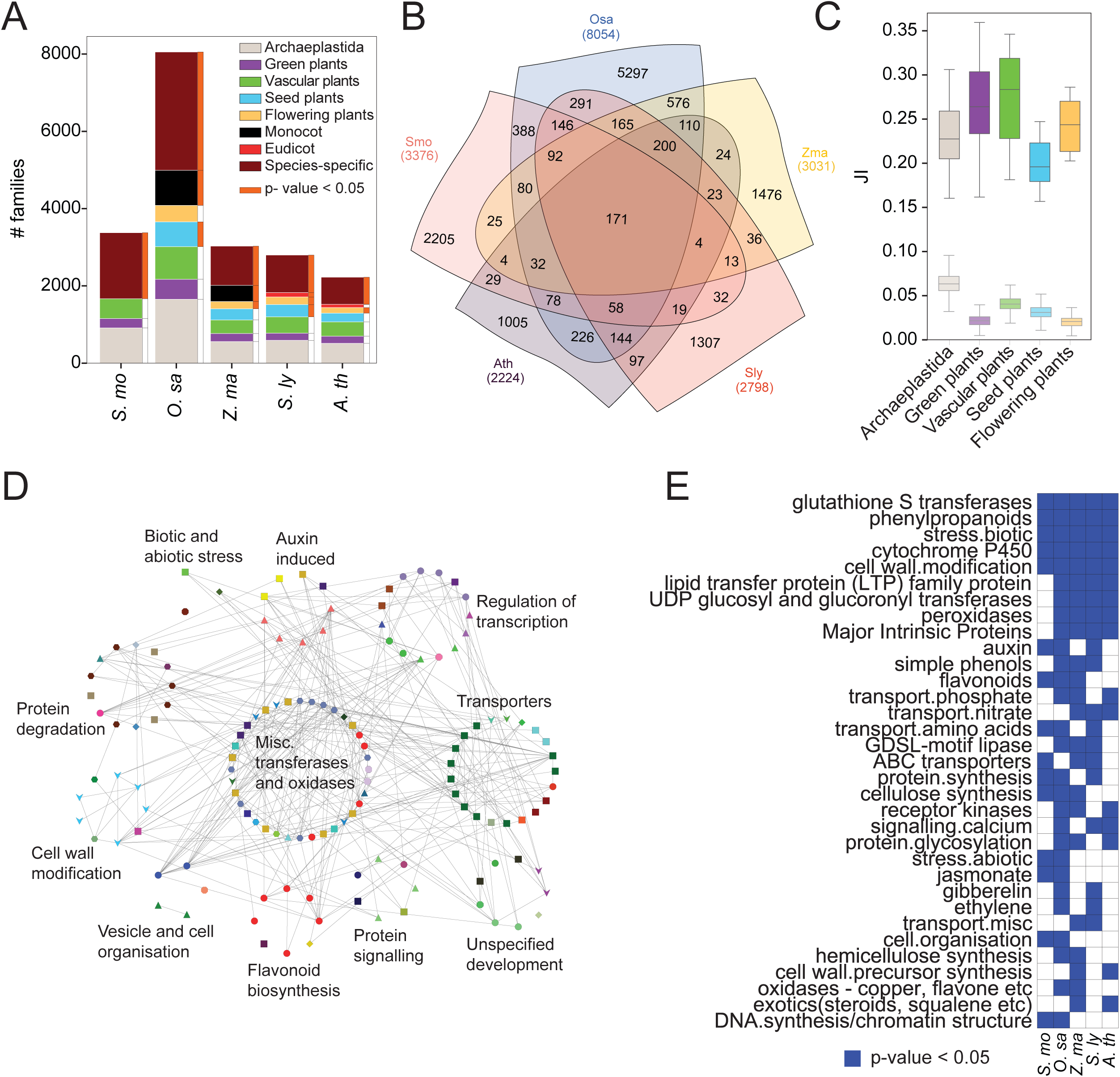
Comparative analysis of the root transcriptomes of *S. moellendorffii* and flowering plants. (A) The number of root-specific gene families whose members are expressed in roots of 5 different plant species. The x-axis indicates the species, while the y-axis shows the number of gene families. The color of the bars indicates the phylostrata of the gene families. (B) Venn diagram showing the overlap of the root-specific gene families in the five plants. (C) The boxplots show the distribution of Jaccard index values (JI, y-axis), capturing the overlap of gene families of genes specifically expressed in roots. The gene families are divided into phylostrata (x-axis), from oldest (Archaeplastida) to youngest (flowering plants). The transparent boxplots (indicated by random) represent the JI values calculated by random sampling of an equal number of genes from a phylostratum. (D) Co-expression network of *S. moellendorffii* genes that belong to a family that contains members specifically expressed in roots in the five analyzed species. The genes (nodes) are grouped according to the biological function. (E) Blue cells indicate the MapMan bins that are significantly enriched (FDR adjusted p-value < 0.05) among the root-specific genes. Bins that are enriched in at least two species are shown.

Next, we investigated the similarities of the root transcriptomes in the five vascular plants by analyzing the overlap of the root-specific gene families. While the majority of the families are species-specific, a considerable amount of families are shared across all five vascular plants (171 families), flowering plants (200 families), monocots (576 families) and eudicots (97 families), suggesting a further elaboration of the root transcriptome in the different lineages (Figure 6B). We then investigated how the different phylostrata contribute to the similarities of the root transcriptomes by calculating the Jaccard Index (JI) between the five phylostrata in all five species pairs. The JI ranges from 0 (no gene families assigned to a given phylostratum are in common) to 1 (all gene families are in common). Despite the differences in the JI values across the phylostrata, we observed that all five phylostrata contributed to the similarity of the root transcriptomes (Figure 6C), and were larger than expected by chance (FDR adjusted p-value < 0.05, Figure 6C, transparent box plots). This further supports the observation that the evolution of roots did not coincide with a punctuated appearance of novel genetic material.

To investigate the role of the core 171 gene families that are root-specific in all vascular plants (Supplemental Table 6), we extracted *S. moellendorffii* root-specific genes belonging to these families and used CoNekT-Plants to create a custom network (https://conekt.sbs.ntu.edu.sg/custom_network/). Functional analysis of this network revealed groups of genes involved in root biology, such as transferases and oxidases ^84–86^, transporters to mediate root signalling and nutrient uptake ^87,88^, auxin-induced genes important for lateral root growth ^89,90^, regulation of transcription, protein degradation, cell wall modification, and flavonoid biosynthesis. Functional enrichment analysis of the root-specific genes in all five species revealed bins corresponding to glutathione S transferase, phenylpropanoids, biotic stress, cytochrome P450, various transporters, cell wall modification, hormone metabolism, and others (Figure 6E). Thus, these genes and gene families constitute prime candidates to study the root molecular biology in vascular plants.

## Discussion

*Selaginella moellendorffii*, a model organism for lycophytes and one of the oldest extant vascular plants, is invaluable to understand the evolution of land plants and vascular plants. In this paper we produced and collected expression data from several organs, tissue types and environmental stresses to expand the eFP Browser (http://bar.utoronto.ca/efp_selaginella/cgi-bin/efpWeb.cgi, Figure 1G) and the CoNekT-Plants database (conekt.plant.tools, Figure 1H) with data from this species. This comprehensive expression atlas can be used to predict gene function and perform comparative analyses among *S. moellendorffii* and the other eight species representing different families in the database.

We first investigated the conservation of the gene module involved in the biosynthesis of lignocellulose in *S. moellendorffii*. We observed that the module contains genes involved in phenylalanine, lignin and cellulose biosynthesis, various polysaccharide-acting enzymes, cell wall structural proteins and MYB transcription factor putatively regulating the expression of these genes (Figure 2)^91^. Importantly, a similar module has been observed in monocots and dicots ^41^, indicating that the lignocellulosic module is at least 400 million years old and most likely present in similar form in all vascular plants. As second-generation biofuels from lignocellulosic biomass are based on the products of this module ^92^, a better understanding of the genes found in this module, their products, and the underlying regulatory networks will allow improvement of lignocellulose in all crops.

All angiosperms possess primary cell walls (PCW), secondary cell walls (SCW), tip growing pollen tubes, and root hairs. While Arabidopsis contains four gene modules dedicated to each cell wall type, the phylogenetic and expression analysis of cell wall-related genes revealed that *S. moellendorffii* contains one gene module biosynthesizing PCW and SCW (Figure 3). This supports the previous analysis suggesting that the diversification of *CesAs* for primary and secondary cell wall was not required for the evolution of vascular tissue ^93^. The duplication and subfunctionalization events of the CesA, COBRA and CslD families in the ancestor of seed and flowering plants produced multiple, non-redundant copies of each gene. What purpose are these gene and gene module duplications serving? Since lycophytes were tall, fast-growing trees in the Carboniferous period, the increased number of cell wall modules in seed plants was not likely important for increasing the production of lignocellulose. Perhaps the duplication of the cell wall module relieved the multi-tasking strain (adaptive conflict) of the ancestral Selaginella module and allowed a further specialization and improvement of PCW and SCW in seed plants. Alternatively, these duplications may serve no purpose beyond duplicating these genes. For example, *CslD2* and *CslD3* are a product of Arabidopsis-specific duplication (Supplemental Figure 4), and both are needed to produce normal root hairs ^94^, indicating that these genes underwent a duplication-degeneration-complementation-type (DCC) functional divergence ^95^. Since gene modules are composed of genes ^59^, it is likely that the same duplication types that govern genes (e.g. DDC, specialization, neofunctionalization, gene sharing) can govern duplications of gene modules.

We observed a linear relationship between the number of genes assigned to a process and the number of clusters enriched for the process (Figure 4D). This relationship is sufficient to explain the observed increase of clusters enriched for processes such as cell wall, secondary metabolism, hormone metabolisms, and miscellaneous in the vascular plants (Figure 4B). Interestingly, we observed clusters enriched for e.g., hormone metabolism in the two algae *C. paradoxa* and *C. reinhardtii*. While these single-cellular organisms do not have or require a hormonal system, they might contain precursors for these pathways. For example, enzymes involved in the biosynthesis of lignin precursors are found in *C. paradoxa* ^96^, and the complete pathway appeared in the moss *Physcomitrella patens* ^97^. These examples suggest that many of the pathways unique to land plants are likely a result of extension and subfunctionalization of ancient pathways present in algae.

Secondary metabolism dramatically expanded in land plants (Figure 5A-B), raising a question on how these new pathways were integrated into the existing cellular machinery. For example, lignin and cellulose synthesis are coordinated on the level of gene expression (Figure 2) and biosynthesis ^98^. Our analysis revealed that secondary metabolic pathways are often transcriptionally coordinated with relevant pathways (e.g., light stress response - carotenoids, biotic stress - lignin Figure 5D, E), suggesting evolutionarily advantageous wiring of these pathways.

It has been previously shown that the root transcriptome is conserved in vascular plants, indicating a common molecular program used in producing roots ^82^. This conservation is remarkable, as the lycophyte and euphyllophyte lineages are thought to have evolved roots independently, based on fossil evidence showing early euphyllophytes without roots at a time when lycophytes already possessed them ^7,8^. The exact origin of roots is thus still under debate, and the two possible theories are (i) the existence of a primitive root developmental program in the common ancestor of vascular plants or (ii) parallel recruitment of largely similar developmental program independently in the lycophyte and euphyllophyte lineages. Our analysis revealed that the root transcriptome is not enriched for gene families that predate the appearance of roots but rather enriched for gene families that appeared after the roots were established (Figure 6A, C). This suggests that root evolution is an ongoing selective process, which could also explain the large variations in root morphology and architecture observed in different species. Similar analyses for leaves, flowers, and other organs and tissues will shed more light on the question whether the evolution of plant structures followed the same trajectory as in roots.

## Methods

### Experimental growth conditions and sampling

*Selaginella moellendorffii* was obtained from Evermoss (http://www.evermoss.de). The plants used for the organ expression atlas and temperature stresses were grown at 24°C under 16 hour light (90 μmol photons m^-2^ s^-1^) and 8 hour darkness photoperiod on soil, while the plants used for high light and darkness stresses were grown on modified Hoagland solution (0.063□mM FeEDTA; 0.5□mM KH_2_PO_4_; 2.5□mM KNO_3_; 2□mM Ca(NO_3_)_2_□×□4H_2_O; 1□mM MgSO_4_□×□7H_2_O; 50.14□µM H_3_BO_3_; 9.25□µM MnCl2□×□4H_2_O; 1□µM ZnCl_2_; 1□µM CuCl_2_; 0.5□µM Na_2_MoO_4_□×□2H_2_O, pH 5.8, 0.8% agar) at 24°C under 12 hour light (70 μmol photons m^-2^ s^-1^) and 12 hour darkness before being subjected to the prolonged darkness (48 hours) and high light (150 μmol photons m^-2^ s^-1^) treatments. The sampled organs were flash-frozen in liquid nitrogen and ground with mortar and pestle. RNA was extracted from the samples using Plant RNeasy Plant kit (Qiagen) according to the manufacturer’s instructions. The integrity of the RNA was assessed using RNA Nano Chip on Agilent Bioanalyzer 2100 and all samples showed RNA integrity values higher than the recommended thresholds. cDNA libraries were prepared from total RNA using poly-A enrichment and sequenced using Illumina-HiSeq2500/4000 at the Beijing Genomics Institute. SEM images were made using freshly harvested and N2-frozen plant samples in a tabletop Hitachi TM3030.

### Estimating gene expression from RNA sequencing data

The reads were mapped to the coding sequences (CDS) with Kallisto ^99^ to obtain transcripts per million (TPM) gene expression values. Publicly available RNA-seq experiments were downloaded from ENA ^100^. The CDSs were downloaded from Phytozome ^1,101^. On average, 55 % of the reads mapped to the coding sequences (Supplemental Table 1). Hierarchical clustering of the samples revealed the expected clustering, where related samples were found on the same clade. The TPM values were combined into a TPM expression matrix (Supplemental Table 2). Due to the generation of a more updated diurnal atlas for *S. moellendorffii* ^102^, the 8 samples generated capturing the diurnal gene expression were excluded from the expression profile but were still used to generate the co-expression network.

### Adding *S. moellendorffii* to the CoNekT-Plants database

The normalized TPM matrix was uploaded in CoNekT-Plants together with CDS sequences ^1,101^, annotation (Mercator, standard settings)^103^ and protein domain information (InterProScan 5.32-71.0) ^104^. The gene families were detected with OrthoFinder 1.1.8 ^105^.

### Functional enrichment analysis of clusters and root transcriptomes

The Heuristic Cluster Chiseling Algorithm (HCCA) clusters were downloaded from CoNekT-Plants (Supplemental Table 3 for Selaginella clusters). For each cluster, we counted the number of times a specific biological process was represented. The observed distribution of biological processes was compared to a permuted distribution obtained by sampling an equal number of genes from the total pool of a species genes for 10,000 permutations. The empirical p-values obtained were false-discovery rate (FDR) corrected by the Benjamini-Hochberg procedure ^106^. The functional enrichment of clusters was performed separately for different levels of MapMan bins, starting from a general first level (Figure 4A, B), including specific bins of secondary metabolism (Figure 5A, B), and more specifically till 3rd level (Figure 5D, E).

The functional enrichment of the root transcriptomes was performed by comparing the observed distribution of MapMan bins within the root-specific genes to a random distribution obtained by sampling an equal amount of genes from the total pool of the species for 10000 permutations. The empirical p-values were FDR corrected (Figure 5D, E, Figure 6E).

### Phylostratigraphic analysis of root transcriptomes

The phylostratum of a family was assessed by identifying the oldest clade found in the family ^24^. In order to test whether a specific phylostratum was enriched in the root transcriptome, we randomly sampled (without replacement) the number of observed root-enriched gene families 10,000 times. The empirical p-values were obtained by calculating whether the observed number of gene families for each phylostratum was larger than the number obtained from the 10,000 sampling procedure. The p-values were FDR corrected (Figure 6A, orange bars, Supplemental Table 7).

To calculate whether a given phylostratum is enriched among the conserved families between two species, we used the Jaccard index (JI), that represents the number of shared families between a species pair. For each phylostratum, the observed JI was compared to the JI obtained by shuffling the family-phylostratum assignment 10,000 times (Figure 6C).

## Data availability

The raw sequencing data is available from EBI accession number E-MTAB-8217.

## Supplemental Figures

**Supplemental Figure 1. Scanning electron microscopy of Selaginella organs and tissues.** (A) SEM image of the microphyll vein. The red and green arrows indicate a stoma and an epidermal cell, respectively. (B) A strobilus with one spore (green arrow) and removed sporophylls. (C) Shoot tip with lateral (red) and medial (green) microphylls. (D) Dissected strobili with spore (green arrow) and sporophylls (red). (E) Isolated sporophyll (green) and spore (red arrow).

**Supplemental Figure 2. Sample relationship dendrogram of the 91 samples.** The dendrogram was produced by calculating the euclidean distance between the expression vectors and clustered using single linkage method.

**Supplemental Figure 3. Phylogenetic trees of the *CesA* genes.** The genes from different species are color coded and their expression in different tissues is shown by the heatmap which indicates high (blue) and low (yellow) expression. The species color coding and gene ID patterns are AT#G##### (brown, Arabidopsis), Solyc##g###### (red, tomato), GSVIVT########### (purple, grape), Zm#####d###### (yellow, maize), LOC_Os##g##### (blue, rice), MA_#####g#### (green, spruce), Smo###### (salmon, Selaginella).

**Supplemental Figure 4. Phylogenetic trees of the *CslD* genes.**

**Supplemental Figure 5. Phylogenetic trees of the *CTL* genes.**

**Supplemental Figure 6. Phylogenetic trees of the *KOR* genes.**

**Supplemental Figure 7. Phylogenetic trees of the *CC* genes.**

**Supplemental Figure 8. Phylogenetic trees of the *COBRA* genes.**

**Supplemental Figure 9. Heuristic Cluster Chiseling Algorithm (HCCA) clusters of *S. moellendorffii*.** The columns show cluster ID, while the rows correspond to MapMan bins that represent biological processes. Clusters enriched for a MapMan bin (FDR adjusted p-value < 0.05) are indicated by red cells.

**Supplemental Figure 10. The relative abundance of secondary metabolites gene families.**

## Supplemental Tables

**Supplemental Table 1. Kallisto mapping statistics.** The first column represents the sample ID, the second, third and fourth columns indicate the number of processed reads, mapped reads and the percentage of mapped reads respectively.

**Supplemental Table 2. TPM normalized expression matrix.**

**Supplemental Table 3. HCCA clusters of *S. moellendorffii*.**

**Supplemental Table 4. Correlation analysis of functionally enriched clusters and number of genes assigned to a specific biological process.** The first column indicates the MapMan binID and its description, the second and third column indicate the R-value and p-value, respectively.

**Supplemental Table 5. Association matrix of biological processes.** The first and second columns represent the two MapMan bins which are associated, the third and fourth columns indicate the number of species the association was found and the species code respectively.

**Supplemental Table 6. Gene families specifically expressed in ‘Roots/rhizoids’ in *S. moellendorffii, O. sativa, Z. mays, S. lycopersicum* and *A. thaliana*.**

**Supplemental Table 7. Phylostrata enrichment of root-specific genes.**

