## Supplementary figures and images for "Expression atlas of *Selaginella moellendorffii* provides insights into the evolution of vasculature, secondary metabolism and roots"

### Fig S1

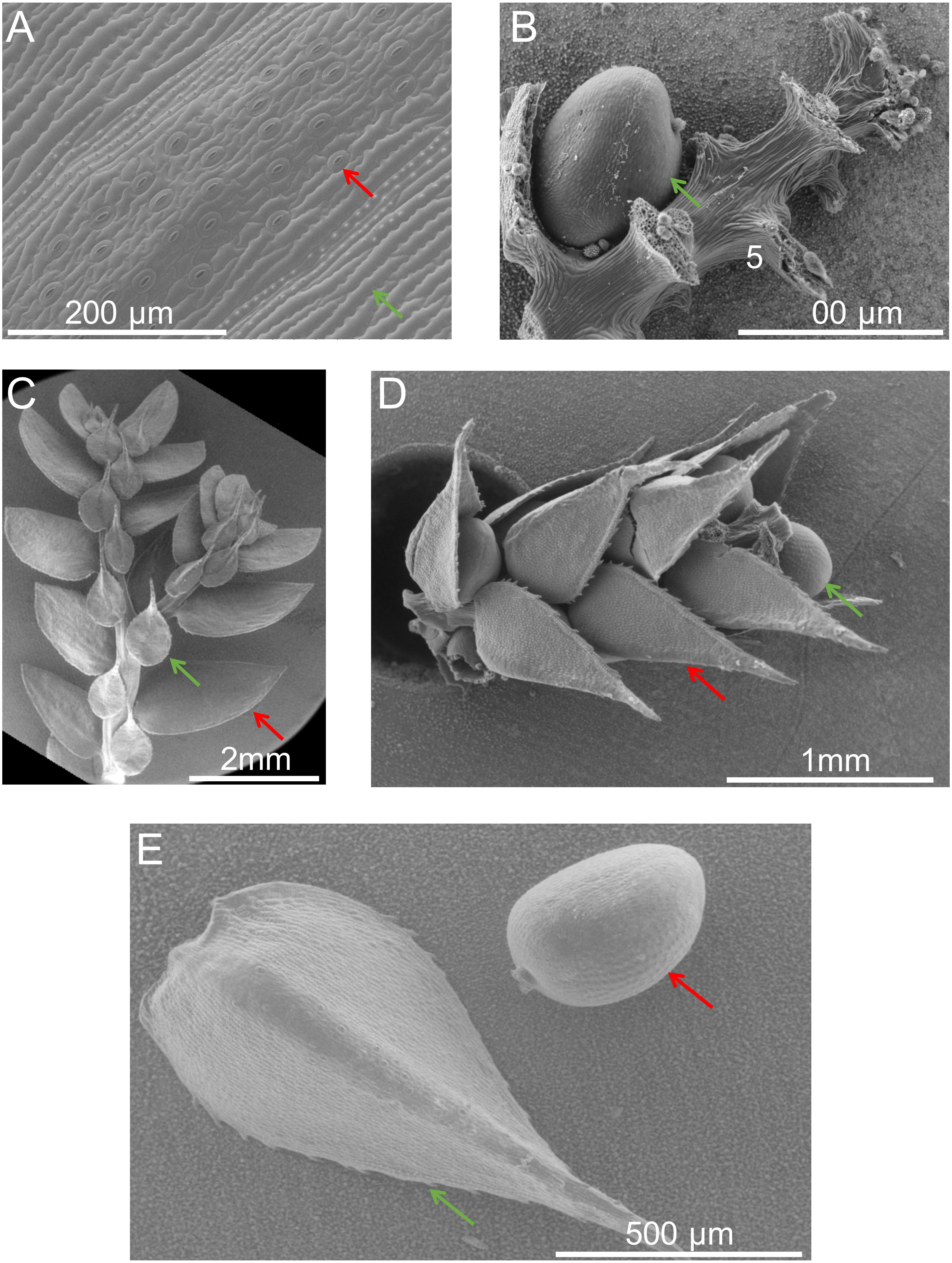

### Fig S2

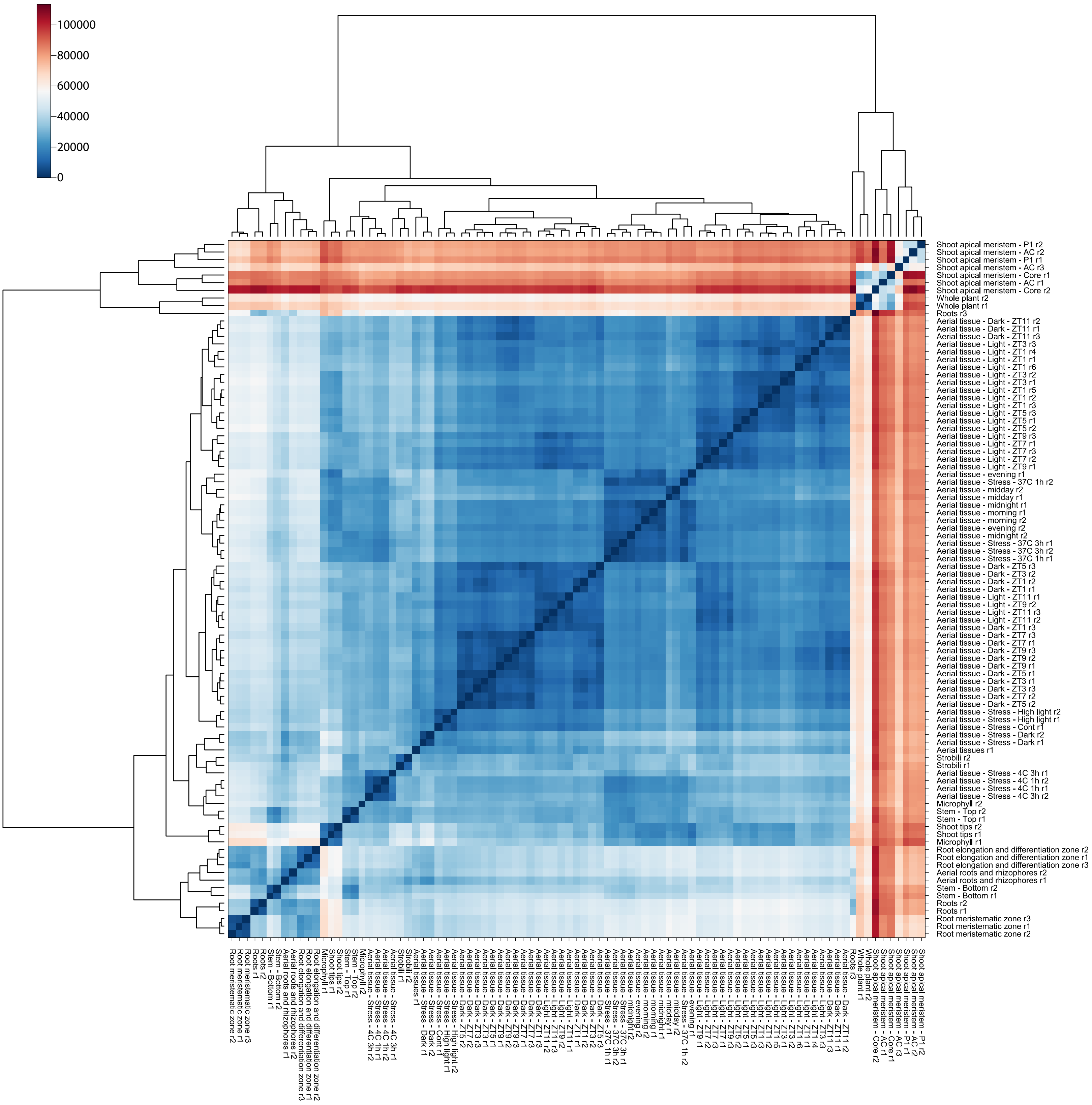

### Fig S3

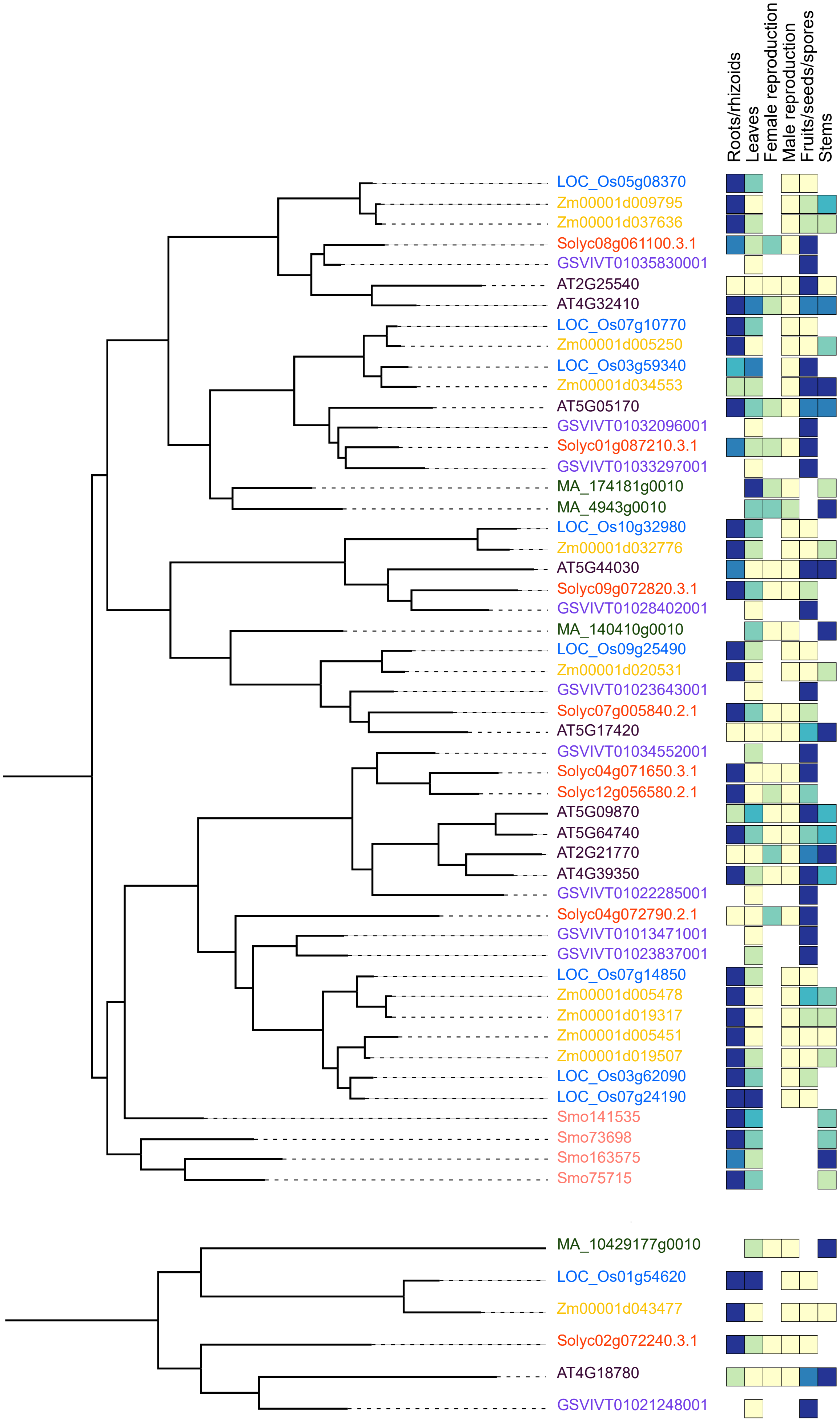

### Fig S4

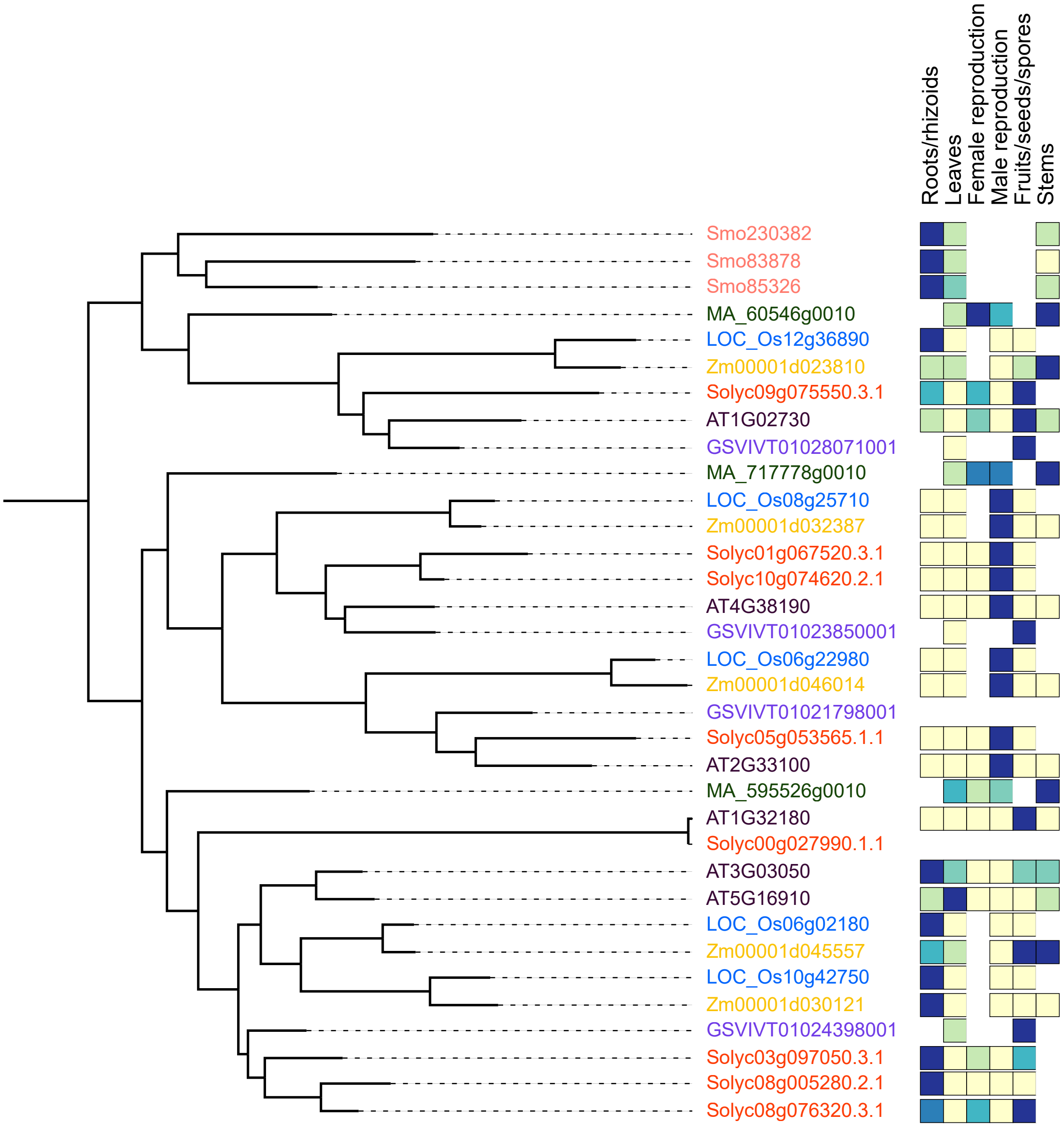

### Fig S5

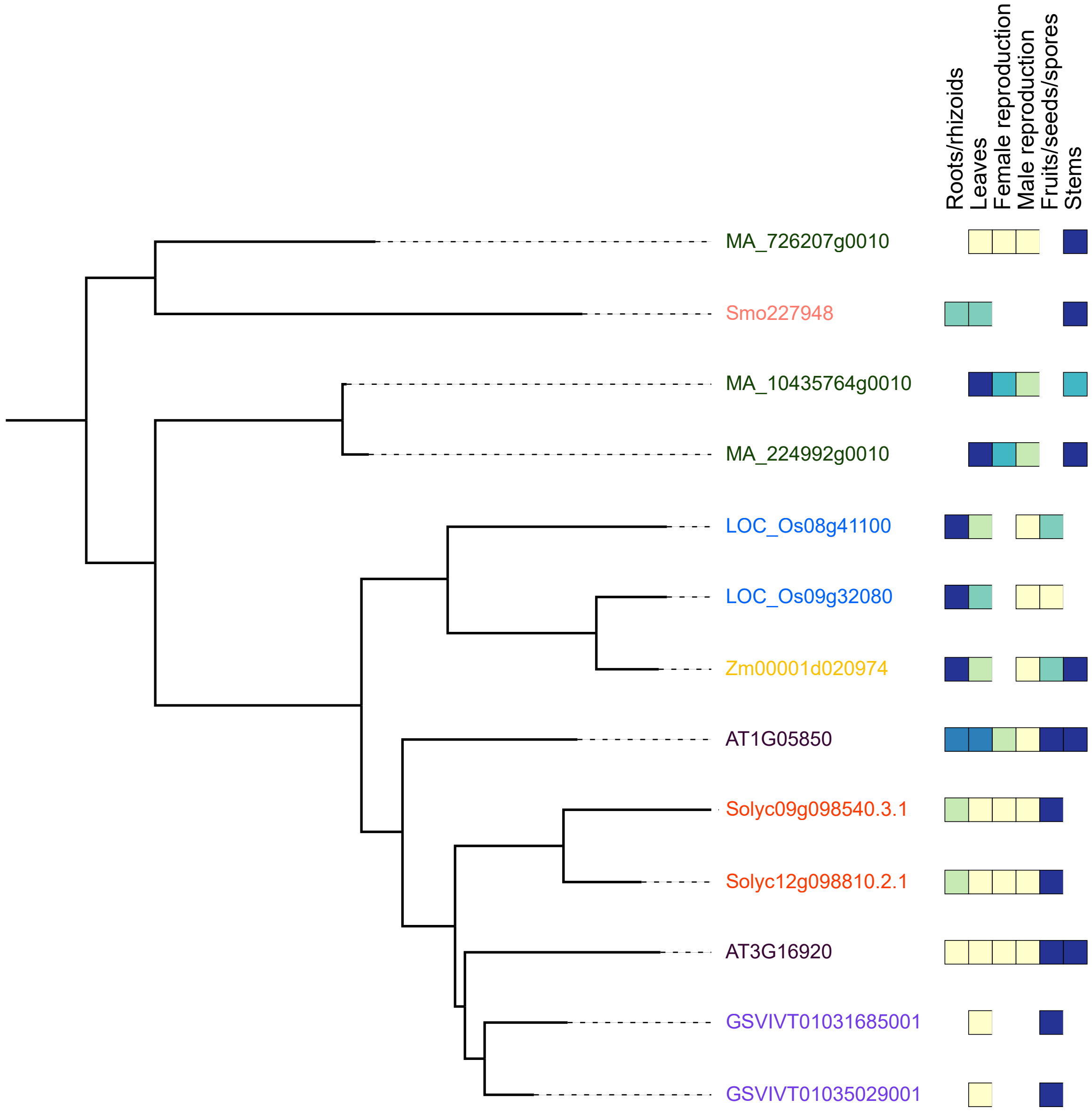

### Fig S6

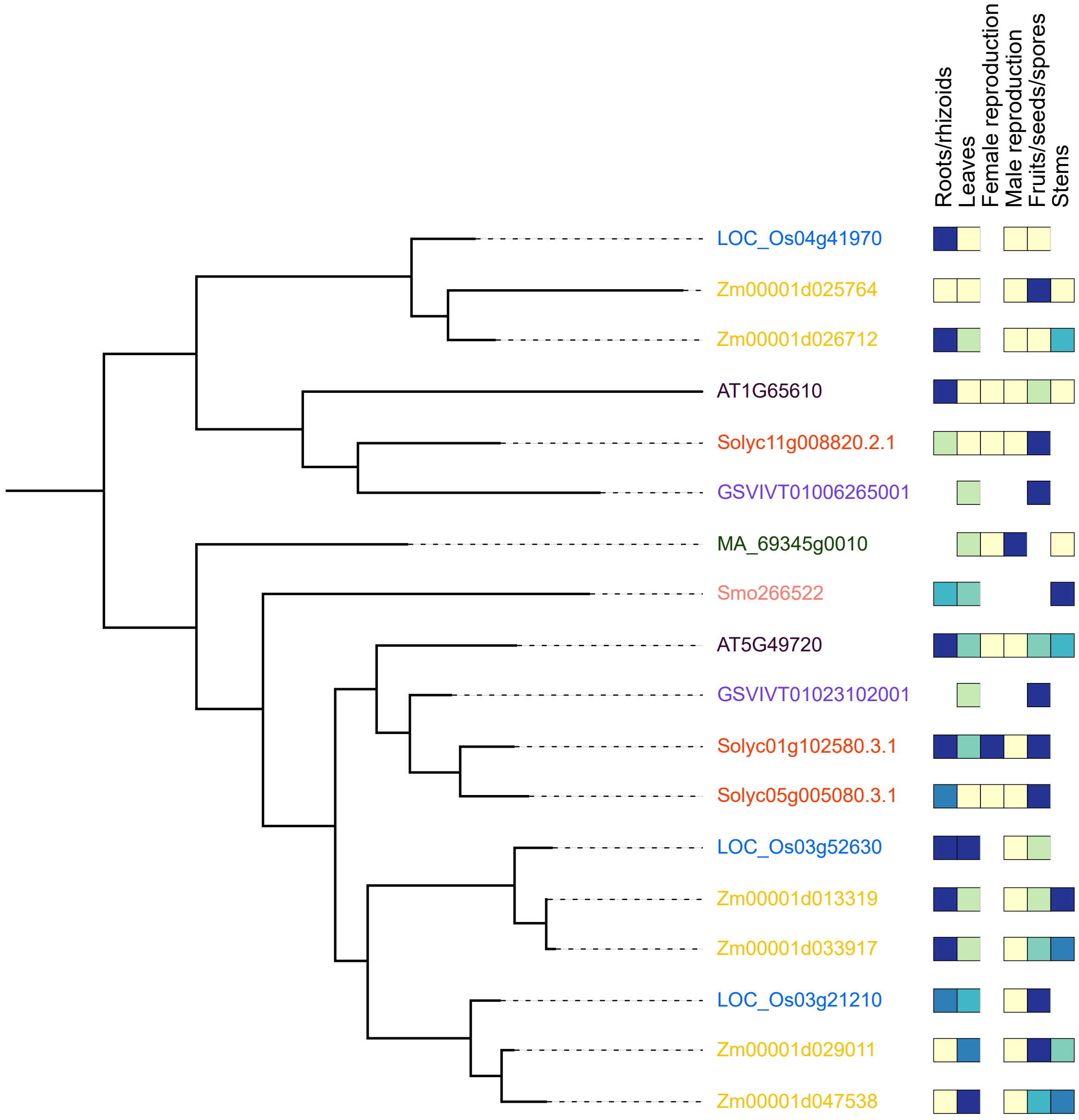

### Fig S7

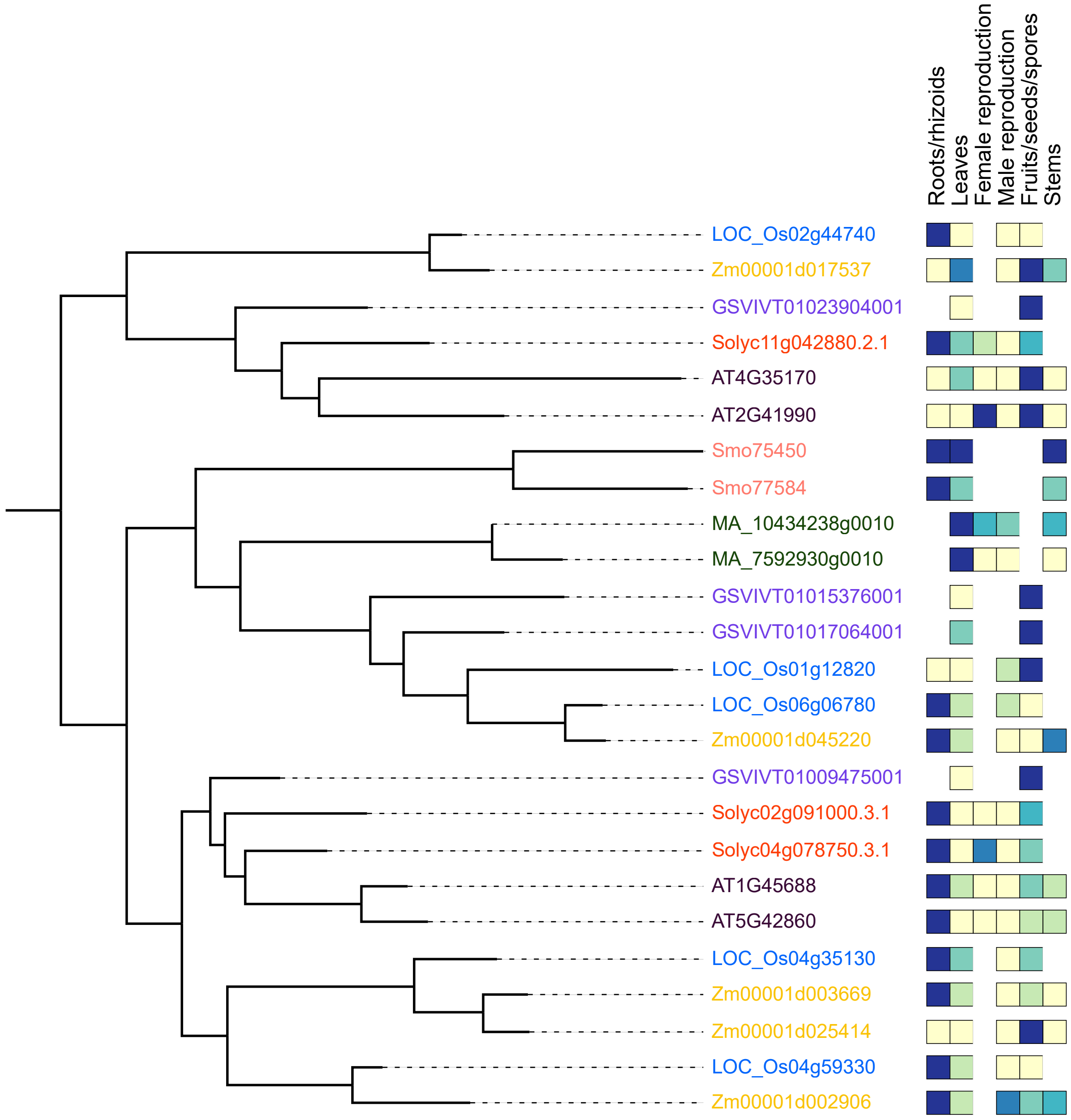

### Fig S8

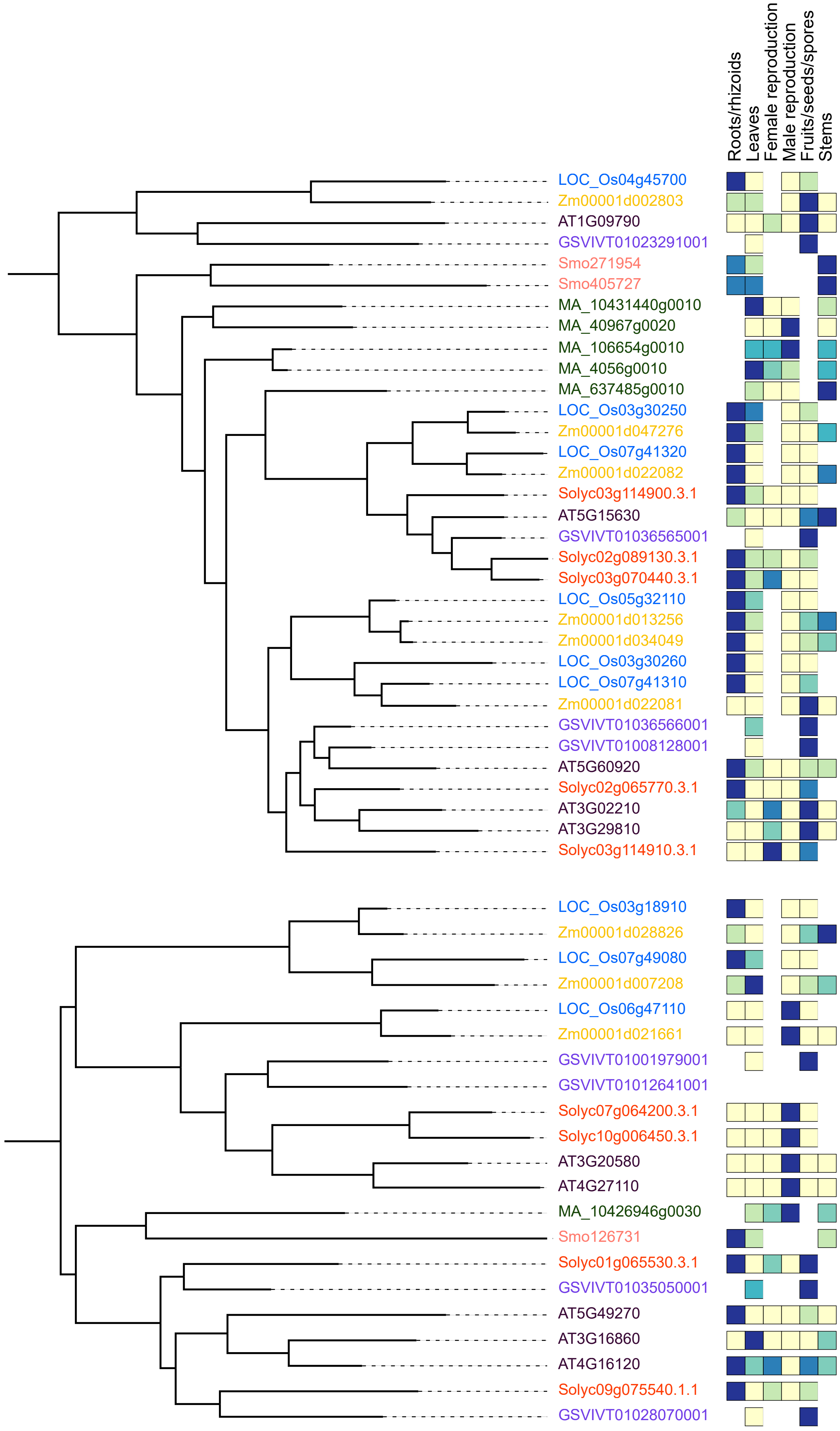

### Fig S9

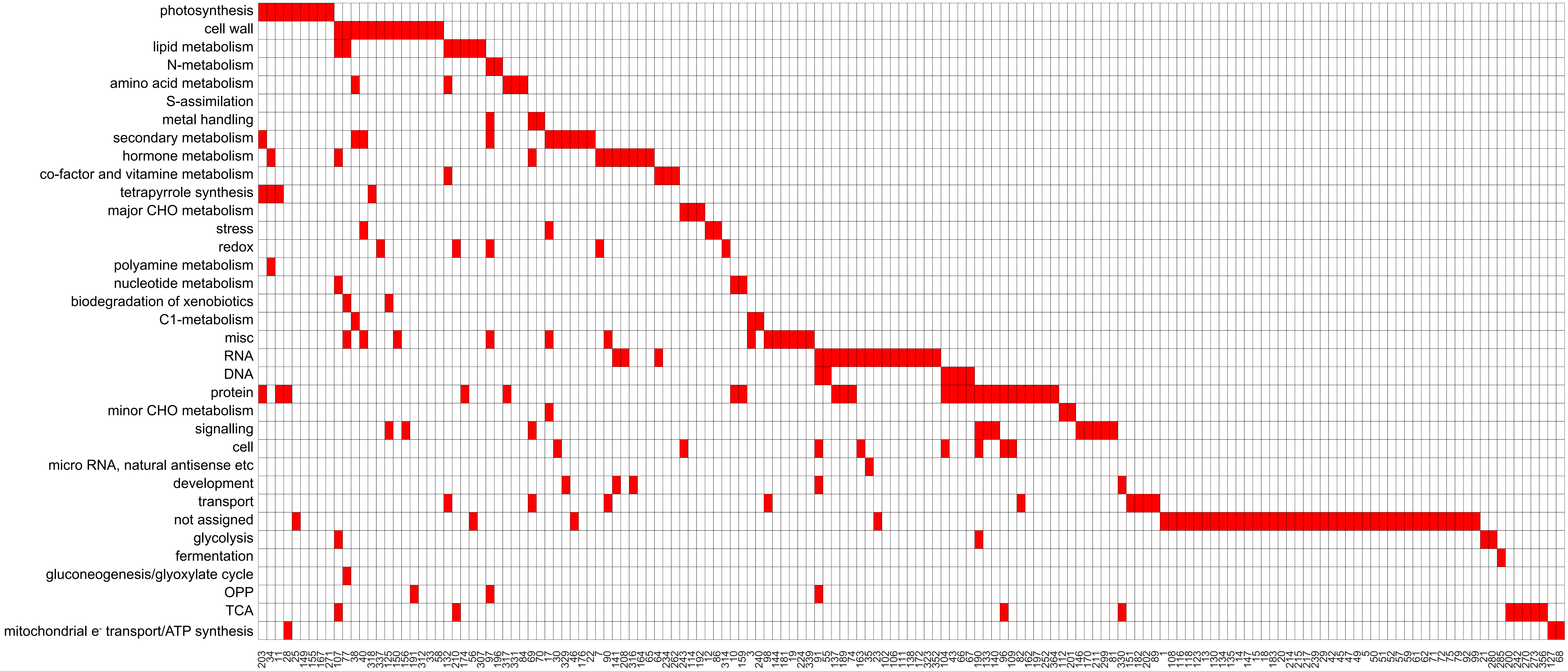

### Fig S10

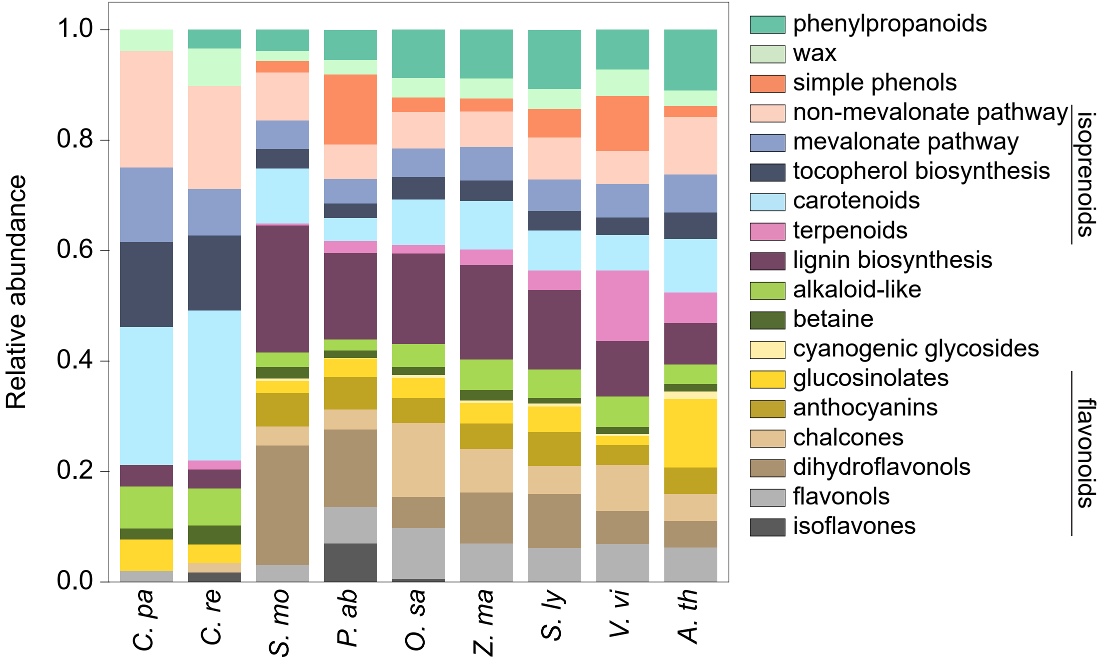
